# A dengue monovalent vaccine with novel structure provides cross-protection against four serotypes of dengue virus

**DOI:** 10.1101/275685

**Authors:** Day-Yu Chao, Wen-Fan Shen, Jedhan Ucat Galula, Jiun-Hung Liu, Mei-Ying Liao, Chang-Hao Huang, Yu-Chun Wang, Han-Chung Wu, Jian-Jong Liang, Yi-Ling Lin, Matthew T. Whitney, Gwong-Jen J. Chang, Sheng-Ren Chen, Shang-Rung Wu

## Abstract

Dengue fever is caused by four different serotypes of dengue virus (DENV) which is the leading cause of worldwide arboviral diseases in humans. The vaccine candidates under development require a tetravalent immunogen to induce a balanced immunity against all four serotypes of dengue virus. Herein we show that mice vaccinated with highly matured virus-like particles derived from DENV serotype 2 (mD2VLP) can generate higher and broader neutralization antibodies (NtAbs) against all 4 serotypes of DENV through clonal expansion supported by hybridoma and B-cell repertoire analysis. The cryo-electron microscopy reconstruction showed that mD2VLP particles possess a T=1 icosahedral symmetry with a groove located within the E-protein dimers near the 2-fold vertices that exposed highly overlapping, cryptic neutralizing epitopes. Most importantly, maternally transferred antibodies derived from mD2VLP-vaccinated female mice protected suckling mice from lethal challenge by all four serotypes of DENV. Our results support the fact that a universal dengue vaccine that protects against all four serotypes of dengue viruses can be achieved by using an immunogen such as mD2VLP.

---

Dengue virus (DENV), a member of the family *Flaviviridae*, is a mosquito-borne pathogen with four distinct serotypes, DENV-1 to DENV-4^1^. It has been estimated that DENV infects about 390 million individuals globally each year, resulting in 96 million clinically apparent infections ranging from mild fever to dengue hemorrhagic fever (DHF) or the life-threatening dengue shock syndrome (DSS)^2^. Although the chimeric yellow fever 17D-derived, tetravalent dengue vaccine (CYD-TDV) has recently been approved by the governments of a few DENV-circulating countries, the unexpected low vaccine efficacy in dengue-naïve children or children younger than 6 years old has limited the use of this vaccine in the 9-45 age group living in endemic countries^3^. Improving the efficacy of CYD-TDV or developing second-generation vaccine candidates is essential to broaden the coverage for all vulnerable populations^4, 5, 6^.

The envelope (E) protein of DENV on the surface of viral particles is the major target of neutralizing antibodies (NtAbs)^7, 8^. The ectodomain of the E protein contains three distinct domains, EDI, EDII, and EDIII, which are connected by flexible hinges to allow rearrangement of domains during virus assembly, maturation and infection^9^. However, the immune response from humans who have recovered from primary DENV infections is dominated by cross-reactive (CR), non-NtAbs which recognize pre-membrane (prM) protein^10^. During virus replication, newly synthesized DENVs are assembled as immature particles in the endoplasmic reticulum at neutral pH, followed by translocation through the trans-golgi network and low-pH secretory vesicles^11^. The pr portion of the prM protein, positioned to cover the fusion loop (FL) peptide at the distal end of each E protein, prevents premature fusion during the maturation process^12, 13^. For DENV to become fully infectious, the pr molecule has to be removed by a cellular furin protease during the egress process^14^. However, furin cleavage in the low-pH secretory vesicles is thought to be inefficient; hence, DENV particles released from infected cells are heterogeneous populations with different degrees of cleavage and release of the pr portion of prM to form the mature membrane (M) protein^15^. Immature DENV (imDENV) particles consist of uncleaved prM protein and partially immature particles (piDENV) containing both prM and M proteins, while fully mature DENV (mDENV) contains only M protein^15, 16^. Functional analyses have revealed that a completely immature flavivirus lacks the ability to infect cells unless in the presence of anti-prM antibodies through a mechanism called antibody-dependent enhancement (ADE) of infection^10, 14^. ADE plays an important role in dengue pathogenesis, and is modulated by the antibody concentration and the degree of virion maturity as shown for West Nile virus (WNV) infection^17^.

Recent studies have suggested that human monoclonal antibodies (MAbs) reactive with all dengue serotypes can neutralize DENV in the low picomolar range. These MAbs have a preference to bind at the envelope dimer epitopes preserved on virion particles with a high degree of maturity^18, 19^. Virus-like particles (VLPs) containing flavivirus prM/E proteins have been demonstrated to be a potential vaccine candidate, since their ordered E structures are similar to those on the virion surface and also undergo low-pH-induced rearrangements and membrane fusion similar to viral particles^20^. Also, VLP vaccines present several advantages since they are highly immunogenic, non-infectious, and accessible to quality control as well as increased production capacity. In this study, we evaluated a mature form of a monovalent dengue VLP for its ability to induce an enhanced immune response with broad neutralizing activity against both homologous and heterologous dengue viruses from all four serotypes.

## Results

We theorized that the process of VLP maturation might be similar to that of dengue viral particles so that VLP antigens would also induce prM-recognizing non-NtAbs. To overcome this potential problem, we engineered the DENV-2 VLPs as completely mature particles by manipulating the furin cleavage site at the junction of pr and M in a DENV-2 VLP-expressing plasmid^21^. For comparison, we also generated imD2VLPs by mutating the minimal furin cleavage motif REKR to REST^12^, which resulted in completely uncleaved prM as detected by western blot and ELISA (Fig. 1). Generation of mature DENV-2 VLP (mD2VLP) was much more challenging. First, we performed multiple sequence alignments of the pr/M junction from several flaviviruses, including the distance-related cell fusion agent virus (CFAV), and analyzed their furin cleavage potential using the algorithm PiTou 2.0^22^. DENV-3 pr/M junction had the lowest predicted Pitou score of 6.87, while WNV had the highest score of 15.4 (Fig. 1A). Secondly, we determined the cleavage efficiency of D2VLP by replacing P1-8 at the pr/M junction site of DENV-2 with sequences from different flaviviruses. Although the Pitou prediction of the CFAV pr/M junction did not result in a high score for cleavage, replacement with P1-8 of CFAV resulted in the most efficient furin cleavage for D2VLP, with only 31% of prM remaining uncleaved as compared to the 100% uncleaved imD2VLP (Fig. 1A). An additional mutation in the P3 residue from E to S of the CFAV P1-8 construct reduced the prM signaling to 13.7% of that of imD2VLP. We thus named this construct with the replacement of CFAV P1-8 plus the P3 E to S mutation as mD2VLP. The pr/M cleavage of this construct was increased to nearly 90% (Fig. 1B). The mD2VLP and imD2VLP, purified from plasmids transfected culture supernatants, showed consistent conformational integrity as ascertained by similar equilibrium banding profiles in 5-25% sucrose density gradients, as well as comparable particle size distributions as determined by negative staining electron microscopy, similar monoclonal antibody (MAb) binding kinetics, and complex glycosylation on the E proteins (Fig. S1).

**Fig. 1.**
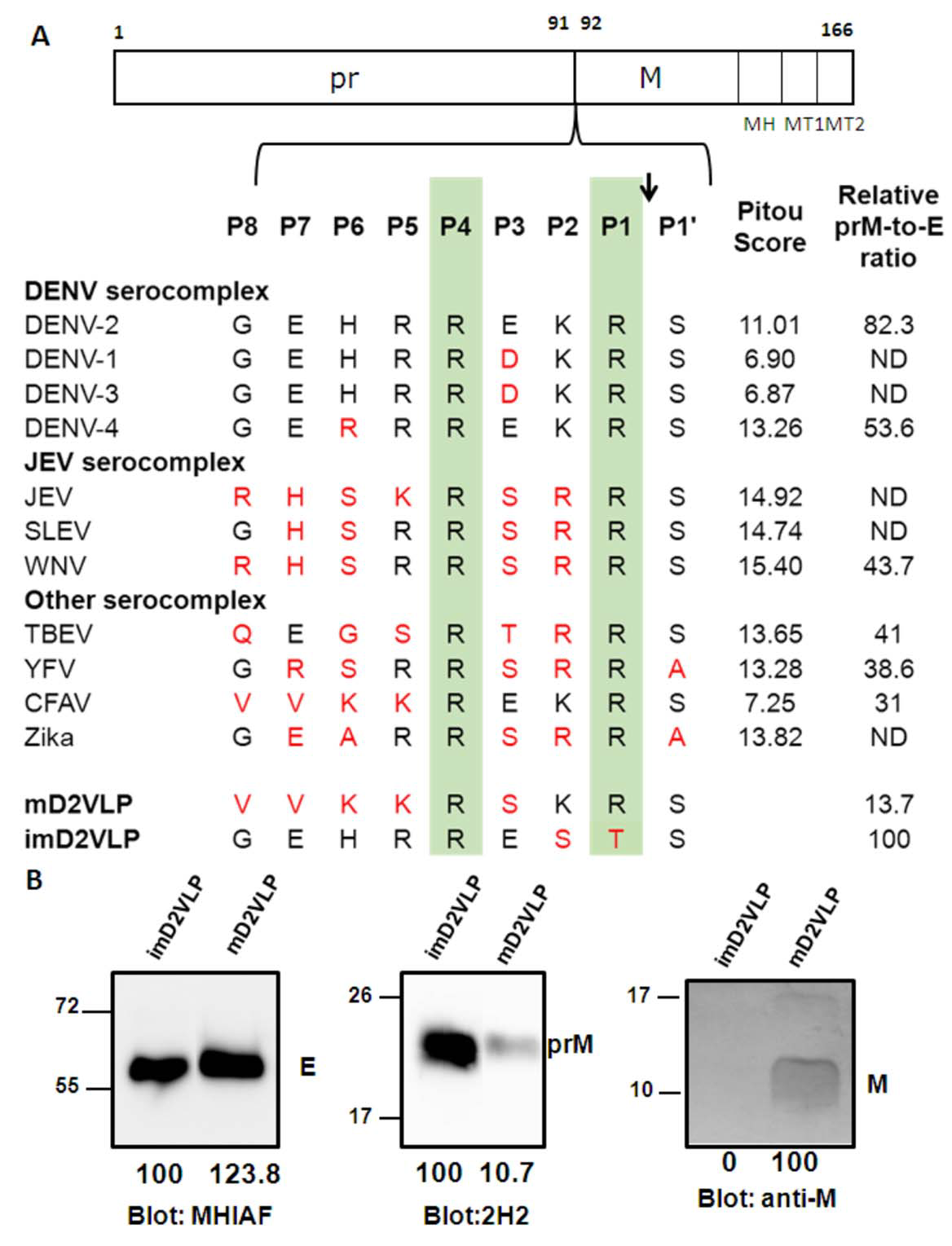
Comparison of the prM junction cleavage efficiency among different DENV-2 virus-like particles (D2VLPs). (A) Schematic drawing of the prM protein. The C-terminal of the prM protein contains an α-helical domain (MH) in the stem region, followed by two transmembrane domains (MT1 and MT2). Numbers refer to the position of the amino acids in the polyprotein starting at the first amino acid of prM according to DENV-2 (NP_056776). Single letter designations of amino acid sequence alignment of representative strains from different serocomplexes of flaviviruses at the prM junction site includes dengue virus serotypes 1-4 (DENV-1 to DENV-4), Japanese encephalitis virus (JEV), St. Louis encephalitis virus (SLEV), West Nile virus (WNV), tick-borne encephalitis virus (TBEV), yellow fever virus (YFV), cell-fusion agent virus (CFAV), Zika virus (ZKV), immature DENV-2 VLP (imD2VLP) and mature DENV-2 VLP (mD2VLP). Numbers with P in the beginning refer to the positions of the amino acid relative to the prM cleavage site in the proximal direction (without apostrophe) and the distal direction (with apostrophe). Protein sequences were aligned, with the key P1 and P4 positions within the furin cleavage sites highlighted. The arrow indicates the prM cleavage site. The amino acids in red indicate the residues different from DENV-2. The PiTou 2.0 furin cleavage prediction scores are shown on the right for each sequence and the higher scores indicate the higher efficiency of furin cleavage. The relative quantity of prM and E of the wild-type and mutant DENV-2 VLP with P1-8 replacement (from other flaviviruses as shown) was measured by ELISA using MAb 3H5 (specific to E domain III of DENV-2) and MAb 155-49 (specific to DENV prM). The relative prM-to-E ratios were calculated by absorbance for prM/absorbance for E protein with reference to imD2VLP, whose pr portion was set as 100% uncleaved, as shown on the right. ND: not determined. Data are presented as means from three representative ELISA experiments with two replicates. (B) Culture supernatants of mD2VLP and imD2VLPs were collected and purified after electroporation with the respective plasmids. Five micrograms of proteins were loaded onto a 12% non-reducing Tricine-SDS-PAGE. E, prM and M proteins were assayed by Western blot using mouse hyper-immune ascitic fluids (MHIAF, 1:2000), MAb 2H2 (0.5 μg/mL) and anti-M protein mouse sera (1:25), respectively. E and prM proteins were visualized with enhanced chemiluminesence (ECL); however, M protein was visualized by TMB substrate to avoid high background. The number below each blot shows the relative densitometric quantification of E, prM and M protein bands by Bio-1D software.

To determine the influence of prM cleavage on the immunogenicity of D2VLP, we immunized groups of 4-week-old BALB/c mice with the purified imD2VLP or mD2VLP (4 μg/mouse) twice at a 4-week interval. Serum samples collected 8 weeks after the boost were analyzed by antigen-specific ELISA for antibody response against homologous D2VLP antigens with different maturity profiles. In order to precisely quantify the amount of dengue-specific antibodies, we used the same quantity of purified VLPs in the antigen-capture ELISA (Fig. 2A). ImD2VLP induced similar ELISA titers of DENV-2-specific IgG antibody against mature and immature antigens (1:22,143 vs. 1:12,804 for imD2VLP and mD2VLP, respectively), although slightly higher titers were observed when using homologous immature antigens. However, stronger reactivity against the mature antigen was noted with mD2VLP immunization (1:15,347 vs. 1:6,381 for mD2VLP and imD2VLP, respectively) (Fig. 2B). We also analyzed the neutralizing ability of these sera against the four serotypes of DENV using a 50% antigen focus-reduction micro neutralization test (FRμNT50). The mD2VLP immunization group induced a higher and broader neutralizing antibody response against all 4 serotypes of DENV (FRμNT50 for DENV-1=1:331, DENV-2=1:597, DENV-3=1:70 and DENV-4=1:141), as compared to the imD2VLP vaccinated group (FRμNT50 for DENV-1=1:100, DENV-2=1:207 DENV-3=1:64 and DENV-4=1:76) (Fig. 2C).

**Fig. 2.**
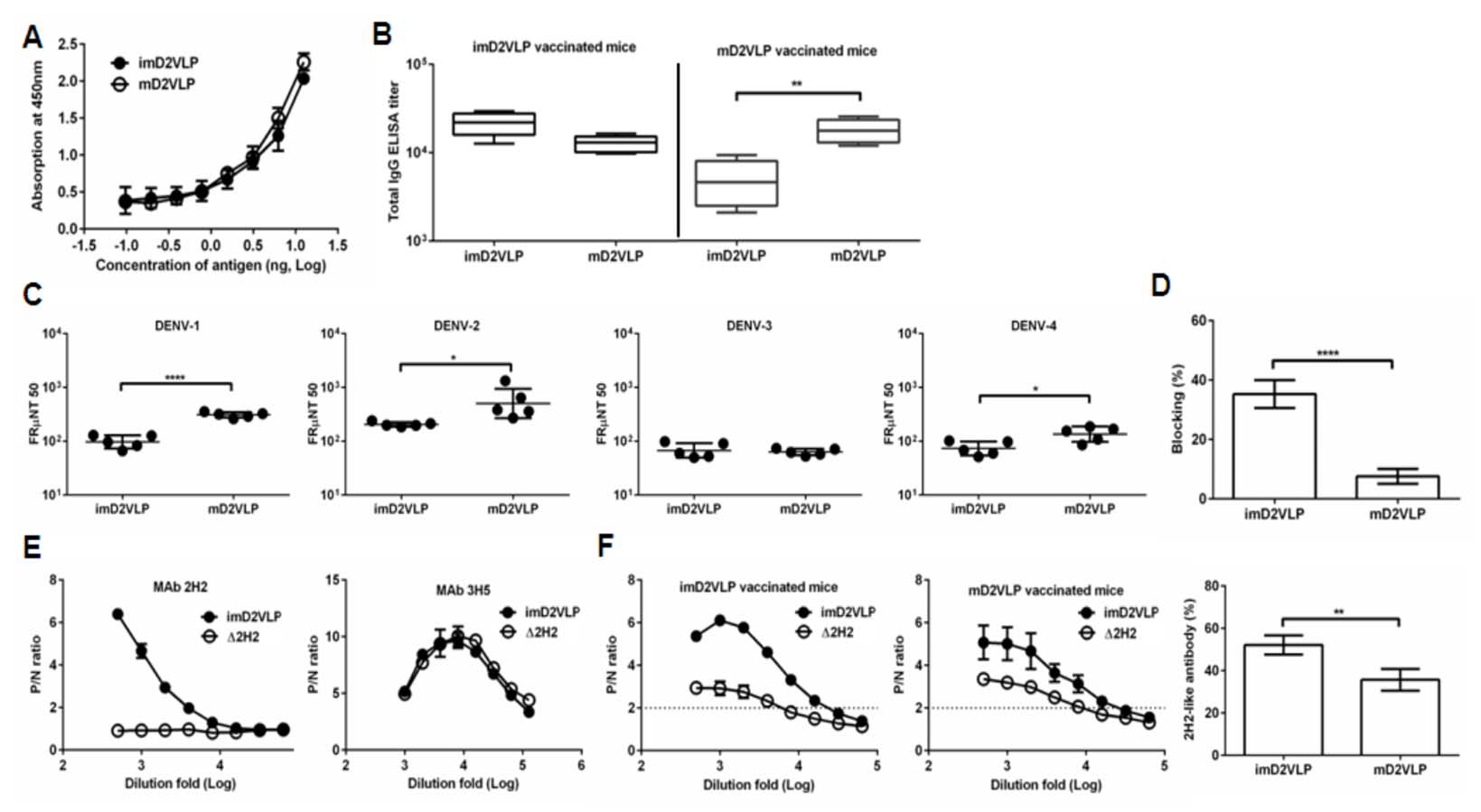
Total antigen-specific IgG, neutralizing titers and proportion of anti-prM antibodies compared between two groups of mice immunized with imD2VLP or mD2VLP. (A) D2VLPs were concentrated and purified from clarified supernatants. The total protein concentration of purified imD2VLP and mD2VLP were first determined by the Bradford assay and then subjected to antigen-capture ELISA using 2-fold serial dilutions. The standard curve was used to titrate both antigens as equal amounts for the subsequent assays. (B) The endpoint IgG titer of 12-week post-immunization mouse sera was measured by antigen-capture ELISA, using equal amounts of homologous and heterologous purified D2VLP antigens. All endpoint titers were log10 transformed and depicted as geometric means with 95% confidence intervals. (C) The neutralizing antibody titers at 50% antigen focus-reduction micro neutralization (FRμNT50) in Vero cells infected with DENV-1 to 4. (D) Proportions of anti-prM antibodies from both D2VLP immunization groups were measured using an epitope-blocking ELISA. Percent blocking of HRP-labeled anti-prM MAb (2H2-HRP) by sera from mice vaccinated with imD2VLP or mD2VLP was determined by the formula 100*[(OD450imD2VLP-OD450imD2VLP blocked by MAb 2H2)/OD450imD2VLP] using a 1:1000 dilution of mouse sera. (E) Serial dilutions of imD2VLP and mutant imD2VLP (Δ2H2), containing mutations at the MAb 2H2 binding site (F1A, K21D, K26P), were tested for binding with MAb 2H2 (left) and control antibody 3H5 (right) by ELISA. P/N ratios refer to the antibody binding magnitude between designated VLP-containing (P) and VLP-free culture supernatant (N) by dividing the absorbance of P by that of N. (F) Binding reactivity of serial dilutions of anti-imD2VLP (left), and anti-mD2VLP (right) mouse sera were analyzed by ELISA using equal amounts of wild-type (imD2VLP) and mutant imD2VLP (Δ2H2) antigens. Proportions of 2H2-like, anti-prM antibodies from two different D2VLP immunization groups were calculated based on the formula 100*[(OD450imD2VLP-OD450Δ2H2)/OD450imD2VLP] at a 1:1000 dilution of mouse sera. All data presented are based on a representative of three independent experiments with two replicates from n=5 mice sera per group per experiment and expressed as mean ± SEM. The statistical significance was determined using the two-tailed Mann-Whitney *U* test to account for non-normality of the transformed data. *, *P* <0.05; **, *P* <0.01;****, *P* <0.0001.

Since the amino acid sequence is identical except for the mutations at the furin cleavage site, the difference in antibody binding and neutralizing activity between mD2VLP and imD2VLP vaccinated mice sera could result from the difference in induction of CR anti-prM non-NtAbs or anti-E NtAbs recognizing structure-dependent epitopes. To address whether the higher neutralization activity induced by mD2VLP was partly due to the reduction of anti-prM antibodies that are known to be cross-reactive but have no neutralizing activity^10^, we performed epitope-blocking ELISAs by using an anti-prM-specific monoclonal antibody (MAb 2H2). MAb 2H2 blocked only 7.59% of the activity of anti-mD2VLP sera but blocked up to 35.26% of antibodies from the imD2VLP-vaccinated groups (Fig. 2D). To avoid steric hindrance due to Mab 2H2 binding, we performed site-directed mutagenesis on three amino acids of pr protein based on the following criteria: (1) the conservation of amino acids among all four serotypes; (2) residues interacting with MAb 2H2^23^; (3) residues not interfering with prM and E interaction^12^. As shown in Fig S2, K26P was a key residue, which significantly decreased the binding of both MAb 2H2 and 155-49 (another cross-reactive murine Mab recognizing pr protein^24^). However, only the mutant Δ2H2-imD2VLP with F1A, K21D and K26P amino acid triple mutations completely the binding of both MAb 2H2 and 155-49 but not that of DENV-2, E-specific Nt-MAb 3H5 (Fig. 2E). The sera from immunized mice were tested for their binding ability to the wild-type and mutant-Δ2H2 imD2VLP. The differences in binding of imD2VLP and Δ2H2 imD2VLP was greater for the anti-imD2VLP sera than the anti-mD2VLP sera (Fig. 2F), suggesting that compared to mD2VLP, imD2VLP was more likely to induce 2H2-like anti-prM antibodies. Thus, mD2VLP has the advantage of eliciting lower levels of anti-prM CR antibody and inducing antibodies with greater neutralizing activity.

To better understand how mD2VLP could induce such a broad antibody response, purified mD2VLPs were subjected to cryo-EM analysis. The results demonstrated that the mD2VLPs had a variable size distribution (Fig. S3). The size variation made improving the resolution challenging, however we were able to determine the overall morphology and E arrangement of mD2VLP. Using a dengue virus-specific Mab (MAb32-6)^25^, immunogold labeling detected the mD2VLPs as spherical particles with a diameter of ~31 nm (Fig. S4), which was also the major population as shown in Fig. S3B. Therefore, we focused on this population for cryo-EM and 3D reconstruction analyses. The results showed that mD2VLPs at a resolution of 15Å had a hollow structure and smooth surface with protrusions around the 5-fold positions (Fig. 3A, left, and Fig. S5). Fitting the atomic E and M surface proteins (PDB: 3J27) into the cryo-EM density map showed that the mD2VLPs followed the T=1 icosahedral symmetry with 30 E dimer subunits on the surface (Fig. 3A, right). The apparent differences of structural features on the mD2VLPs compared to the mature native virion particles^13, 26^ was a slimmer lipid bilayer and the rearrangements of the surface proteins. First, the central section through the viron and m2DVLP reconstructions showed that the bilayer of m2DVLP was relatively thinner than that of viron. The distance between the exterior leaflet and the interior leaflet of m2DVLP and virion was 12.8Å and 14.4Å, respectively (Fig. 3B). A similar T=1 icosahedral symmetry of the E-protein arrangement and thinner lipid bilayer were also found in TBEV VLPs^27^. Second, interpretation of the map showed that E dimer subunits moved apart from each other causing less density in the intra-dimeric interphase (Fig. 4C). Because of this loose interaction, a groove located within the E-protein dimer near the icosahedral 2-fold vertices on mD2VLP was noted (Fig. 4A, right, and Fig. S6). Third, the central section showed that there was a solid density in the H-layer, which is composed of H helices of E and M protein stem regions, while the density in the T-layer, which contains transmembrane helices of E and M proteins, was weaker in the VLP (Fig. 3B). Each E protein consists of an E ecto-domain and a stem region that connects the ecto-domain to its transmembrane region. The stem contains two α-helices, one of which interacts with the viral lipid membrane and the other with the E ecto-domain. This looser interaction of E protein dimers was further stabilized by the E protein rearrangement and the closer interaction with one of the α-helices, which resulted in the solid density beneath the ecto-domain as shown in Fig. 3B. It is worth noting that the stem region of this mD2VLP was replaced into Japanese Encephalitis viral (JEV) sequence; it was previously suggested that this replacement increased the hydrophobicity for the interaction with the lipid membrane and enhanced the secretion of dengue VLP^28^. This rearrangement of E protein under T=1 symmetry on the VLP surface also exposed more accessible epitopes (48.2%; 191 amino acid with ≥30% solvent accessibility among 396 amino acids of the entire E protein) compared to DENV-2 virions (43.4%; 172 amino acid), particularly located at fusion loop, aa 239, aa 251-262 of EDII which together formed groove, and at A strand, cd loop and G strand of EDIII which surrounded a 5-fold axis (Fig. 4A, left, Fig. S7). Footprint analysis (Fig. 4B) showed that those cryptic residues on the E protein, which were previously buried inside of the native DENV virion and only interacted with the broad NtAbs while the virion “breathed”^29^, were better exposed on the m2DVLP. Specifically, the epitopes interacting with MAb 1A1D-2, which binds to the virus at 37 °C30; MAb 2D22, which could block two-thirds of all dimers on the virus surface, depending on the strain^31^; and the ‘E-dimer-dependent epitope’, which is recognized by broadly neutralizing MAbs EDE1 and 219, were all well-exposed in VLPs without steric hindrance.

**Fig. 3.**
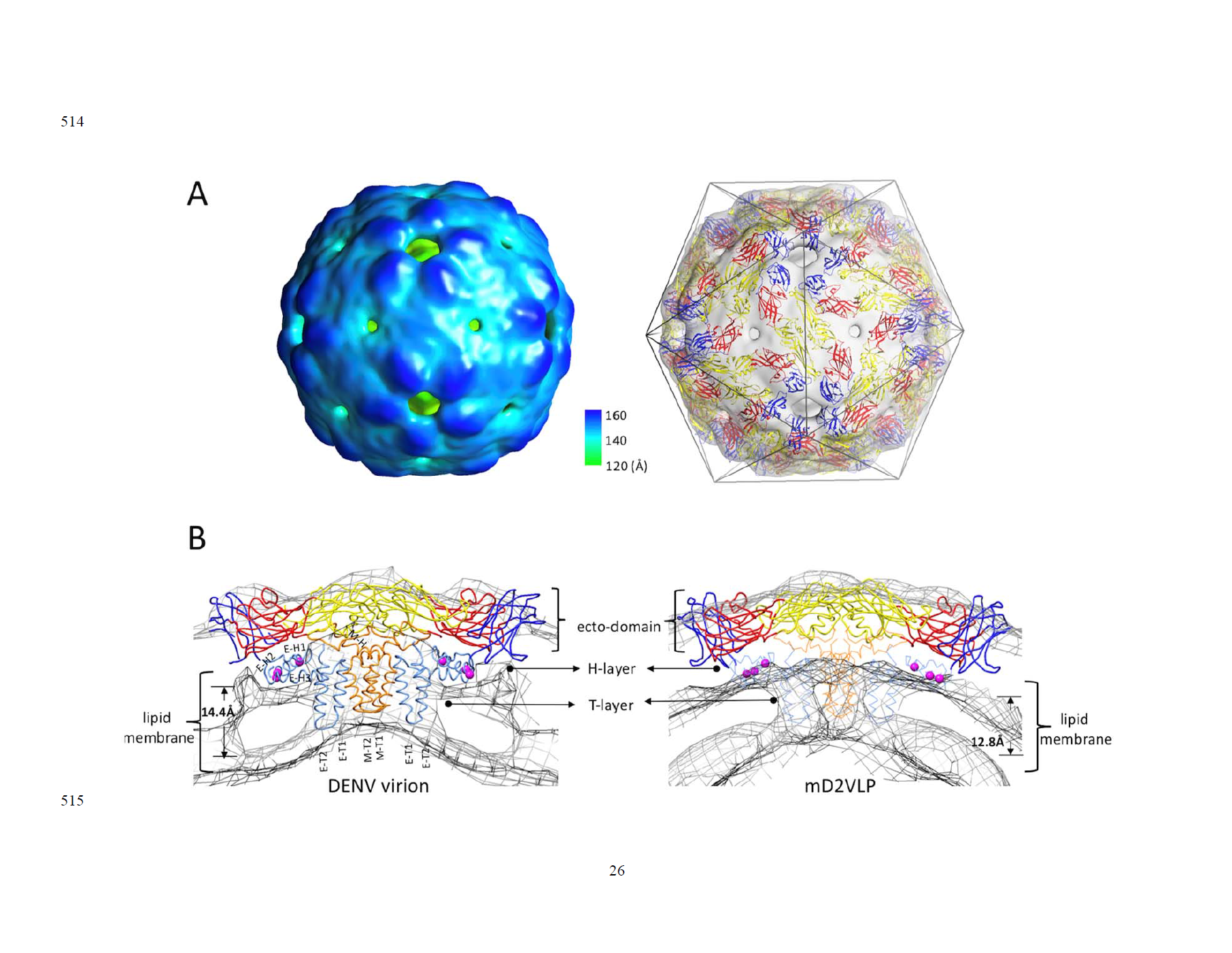
The structure of mature form virus-like particles of dengue virus serotype 2 (mD2VLP). (A) The reconstructed cryo-EM map of the DENV VLPs (left panel) were presented with the radial color-code indicated (left), the size of the particle is 31nm. Fitting of atomic E, M surface protein (PDB: 3J27) into cryo-EM density map (right) showed that the VLP structure followed the T=1 icosahedral symmetry lattice with 30E dimer subunits on the surface. The density map was shown as a transparent volume rendering into which was fitted the backbone structures of the E ecto-domain. The domains I, II, and III were highlighted in red, yellow and blue, respectively. The cage indicated the icosahedral symmetry. (B) The cross section showed the fitted E:M:M:E heterotetramer (PDB: 3J27) into DENV virion (left) and into mD2VLP (right). The map density was in mesh presentation. The atomic model of E:M:M:E heterotetramer was showed in ribbon. Domain definition of dengue E was the same as the previous description, the transmembrane domain of E was colored as light blue, the M protein was in orange color. It was clear that the density of H-layer which is composed of E-H1 to E-H3 and M-H was more solid while the density in T-layer which contains E-T1, E-T2, M-T1 and M-T2 was weak in VLP than in virion. The residues at 398, 401 and 412 in E-H1 of JEV sequence which were proved to play important role in promoting extracellular secretion^28^ were shown as magenta spheres. The transmembrane, perimembrane helices and lipid bilayer were labelled, the critical measurements were also shown.

**Fig. 4.**
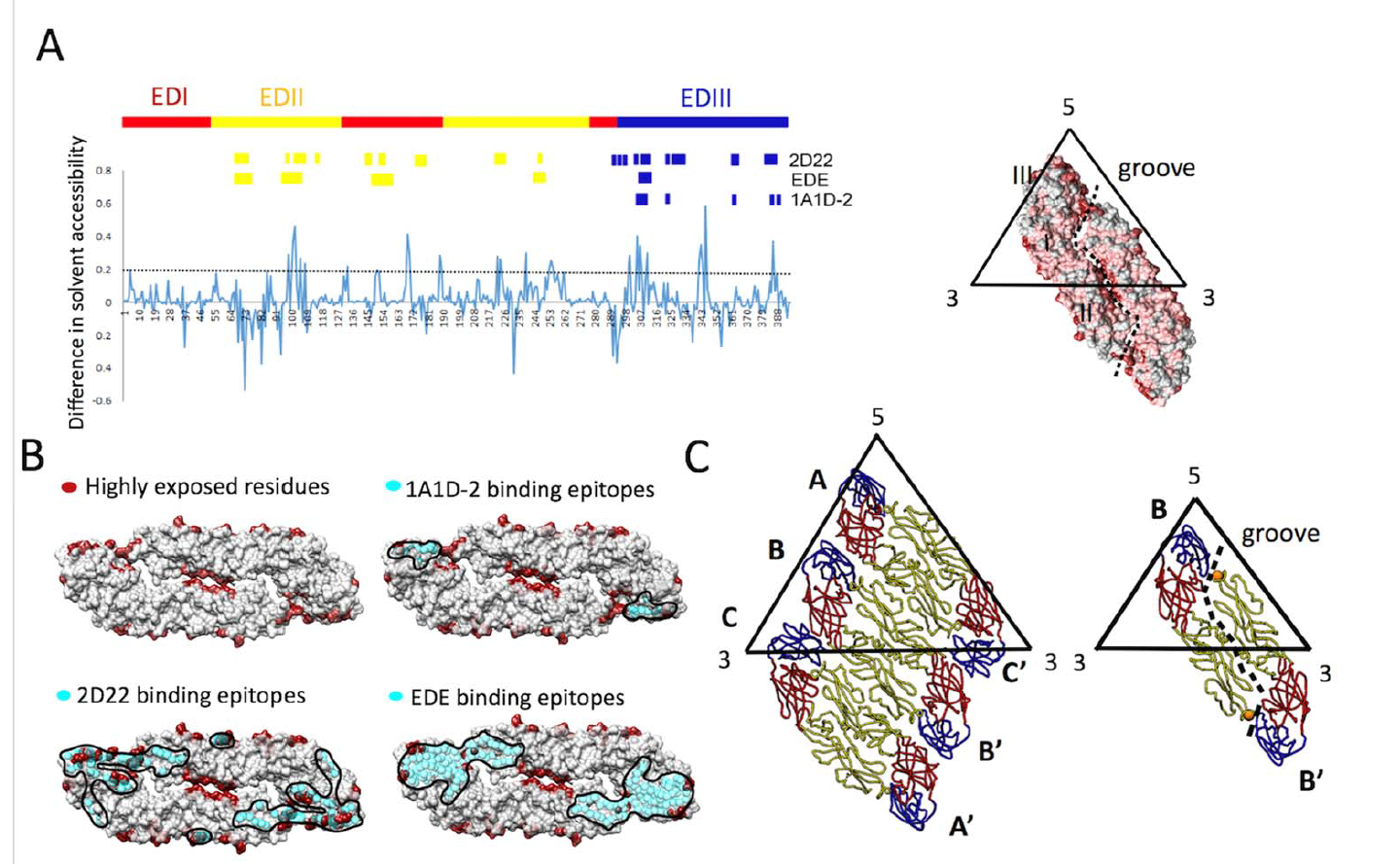
Solvent accessibility of dengue virus serotype 2 virus soluble envelope (sE) protein (A, left) The plot of difference in relative solvent-accessible surface area (Δ%SASA). A positive value of %SASA meant the residue became more exposed when the particle assembly shifts from virion (*T*=3) to VLP (*T*=1). The black dash line indicated the Δ%SASA ≥ 0.2 which was defined as highly exposed residues in VLP comparing to virion. The high positive values which were focused in the peptide regions such as the fusion loop peptide (including amino acid (aa) residues ranging from 100-110), aa 169-170 at domain I, aa 222-226, aa 239 and aa 251-262 at domain II as well as A strand of domain III (aa 300-308), the cd loop of domain III (aa 342-348) and G strand (aa 386-388) around the 5-fold openings. The residues interacting with MAb 1A1D-230, including residues 305-312, 352, 364, 388 and 390; the residues interacting with MAb 2D2231, including residues 67-72, 99, 101-104, 113, 177-180, 225-227, 247, 328, 384-386 (Heavy chain); 148-149, 153-155, 291-293, 295, 298, 299, 307, 309-310, 325, 327, 362-363 (light chain) and the residues interacting with human MAb EDE antibodies^19^, including aa residues 67-74, 97-106, 148-159, 246-249 and 307-314 were indicated. (A, right) The high positive peaks (Δ%SASA ≥ 0.2), low positive peaks (Δ%SASA between 0 and 0.2) and negative peaks (Δ%SASA <0) in the plot were colored by dark red, deem red and grey in the E dimer surface rendering. The groove located within E-dimer interface was outlined. (B) The highly exposed residues (Δ%SASA ≥0.2) which were colored by dark red were shown in the surface rendered E-dimer. The residues in E interacting with MAb 1A1D-2, 2D22 and EDE were in cyan spheres showing that they were highly exposed on the m2DVLP surface. Importantly, the binding footprints of the three antibodies were highly overlapping with footprint of highly exposed residues in m2DVLP, and formed a neutralization sensitive patch on m2DVLPs. The areas of the interacting epitopes are circled by black lines. (C) The E protein forming the rafts in virion were shown in the left panel where the three individual E proteins in the asymmetric unit are labeled A, B, and C of the E proteins, in the neighboring asymmetric unit are labeled A’, B’, and C’. The icosahedral 2-fold E protein dimers (B and B’) in m2DVLP have moved apart from each other causing the groove (right). The aa 101 which was responsible for DM25-3 antibody binding were shown in orange spheres.

The preservation of neutralizing epitopes on the surface of VLPs is critical for efficient production of broadly neutralizing antibody responses. We next investigated if a broad neutralization response to mD2VLP immunization was due to the induction of antibodies with a preference to bind at the E dimer epitopes preserved on the surface of mature VLP^18, 19^. The splenocytes from mD2VLP-vaccinated mice were used to perform fusion and generate hybridomas. Among all hybridomas screened, the reactivities of two representative MAbs were measured by ELISA (Fig. 5A). The results showed that DM8-6 is a serotype-specific MAb reactive only with DENV-2; while MAb DM25-3 recognized all four serotypes of DENV. Next, we tested the neutralizing activity of these two MAbs against four serotypes of DENV. DM8-6 showed good neutralizing activity against DENV-2 at a concentration of 0.037 μg/mL in FRμNT50 but poorly neutralized the other three serotypes. Consistent with the ELISA results, DM25-3 neutralized all four serotypes in FRμNT50 at 0.32, 0.38, 0.24 and 0.58 μg/mL for DENV-1 to DENV-4, respectively (Fig. 5B).

**Fig. 5.**
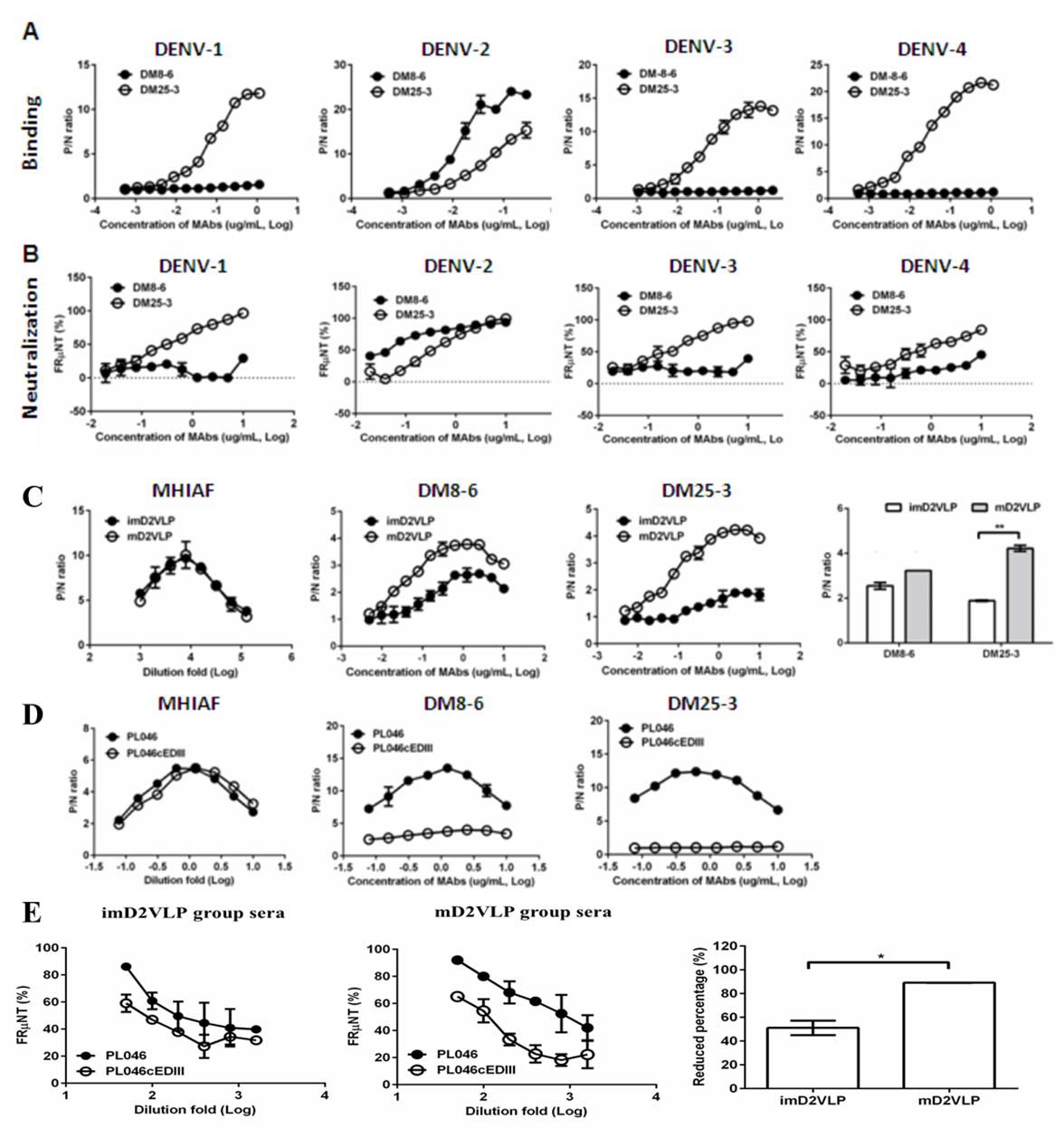
Characterization of murine monoclonal antibodies (MAbs) generated from mouse splenocytes following immunization with mD2VLP. Binding (A) and neutralizing (B) activities of MAbs DM8-6 and DM25-3 against DENV-1 to 4 were measured by ELISA and the focus-reduction micro-neutralization test (FRμNT). (C) Binding curves of MAb DM8-6 (center left) and DM25-3 (center right) against imD2VLP and mD2VLP were performed using ELISAs. Equal amount of both imD2VLP and mD2VLP was properly titrated by antigen-capture ELISA using mouse hyper-immune ascitic fluid (MHIAF) against DENV-2 (left). The difference in binding activities of both MAbs is presented by a bar graph (right). (D) A recombinant DENV-2 virus was produced by replacing domain III with a consensus sequence of domain III (PL046cEDIII)^31^ and the binding activity of DM8-6 (center) and DM25-3 (right) was compared with that of parental DENV-2 strain PL046. Equal amount of both PL046 and PL046cEDIII was properly titrated by antigen-capture ELISA using mouse hyper-immune ascitic fluid (MHIAF) against DENV-2 (left). (E) FRμNT of two-fold diluted mice sera immunized with mD2VLP and imD2VLP against parental PL046 and PL046cEDIII DENV-2 viruses (n=5 per group per experiment). The differences in FRμNT50 from mD2VLP and imD2VLP immunization groups were converted to bar chart at 1:1000 fold dilution of mice sera. The conversion was based on the formula 100*[ FRμNT50 of (PL046- PL046cEDIII)/FRμNT50 of PL046]. P/N ratio refer to the antibody binding magnitude between designated VLP-containing (P) and VLP-free culture supernatant (N) by dividing the absorbance of P by that of N. The data are presented as means ± SEM from three independent experiments with two replicates. The two-tailed Mann-Whitney *U* test was used to test statistical significance. *, *P* <0.05. **, *P* <0.01.

To determine if MAb DM25-3 recognized quaternary structure-dependent epitopes presented only on mature virion particles^19, 31^, DM25-3 was tested for its binding activity to both mD2VLP and imD2VLP using antigen-capture ELISA. MAb DM25-3 recognized mD2VLP very well, but reacted poorly with imD2VLP (Fig. 5C). Next, we performed site-directed mutagenesis on the fusion loop peptide in domain II and A-strand in domain III, both of which are important binding regions for CR group/complex neutralizing antibodies^32^, ^33^ (Fig. S8). The results suggested that amino acid E-101 was likely important in the binding site of DM25-3. Residue 101 on the fusion loop of E protein is conserved among all four serotypes of DENV and can only be exposed when virion particles undergo low-pH induced conformational change or during the “breathing” state^34,35^. Fig. 4C shows that residue 101 is located at the touching point between EDII and EDIII of the dimeric molecule. Since residue 101 is a critical residue in stabilizing E-protein dimers on DENV virion, mutation on 101 would disrupt viral particle formation^26, 36^. However, mutation of residue 101 on mD2VLP can still form particles (Fig S9), which further support our cryoEM model that the E:E protein interactions on VLP are loose. When the E:E protein interaction was looser, residue 101 at the groove within the dimeric molecule was more exposed (64%) than on the native virion (23.8%, PDB:3J27) or on the native virion during the 37oC “breathing” state (45.3%, PDB:3ZKO). We also generated a recombinant virus of DENV-2 whose EDIII was exchanged for a consensus EDIII^37^ and the recombinant virus PL046cEDIII was recovered from the transfected cell culture supernatants for use in ELISA and FRμNT. Compared to the parental DENV-2 strain PL046, there was significant loss of binding of MAb DM25-3 and immune mouse sera to PL046cEDIII (Fig. 5D), suggesting that DM25-3 could be an E-dimer inter-domain antibody whose binding footprint is sensitive to amino acid changes in the EDIII domain. By comparing FRμNT50 tests using murine antisera on both PL046 and PL046cEDIII viruses, we found that anti-mD2VLP sera showed a greater difference in neutralizing activity against PL046 and PL046cEDIII than did anti-imD2VLP sera, indicating a greater conformational dependence for anti-mD2VLP-triggered neutralization (Fig. 5E).

Single B cells from the spleens of mD2VLP or imD2VLP-vaccinated mice were sorted into 96- well plates to better understand DENV-2 prM/E-specific B-cell repertoires (Fig. 6A). By analyzing the amino acid sequences of the RT-PCR products of the variable regions of Ig heavy-chain (IgH) and light-chain (IgL) genes, the B-cell response from mD2VLP-immunized mice was more complex than the imD2VLP-immunized animals, particularly the gene loci of IgH (Fig. 6B). The IgH and IgL genes from the DM25-3-producing hybridoma were also analyzed and found to contain IgHV1-22*01 and IgKV3-12*01 (Fig. S10), which were clonally expanded as shown in Fig. 6B.

**Fig 6.**
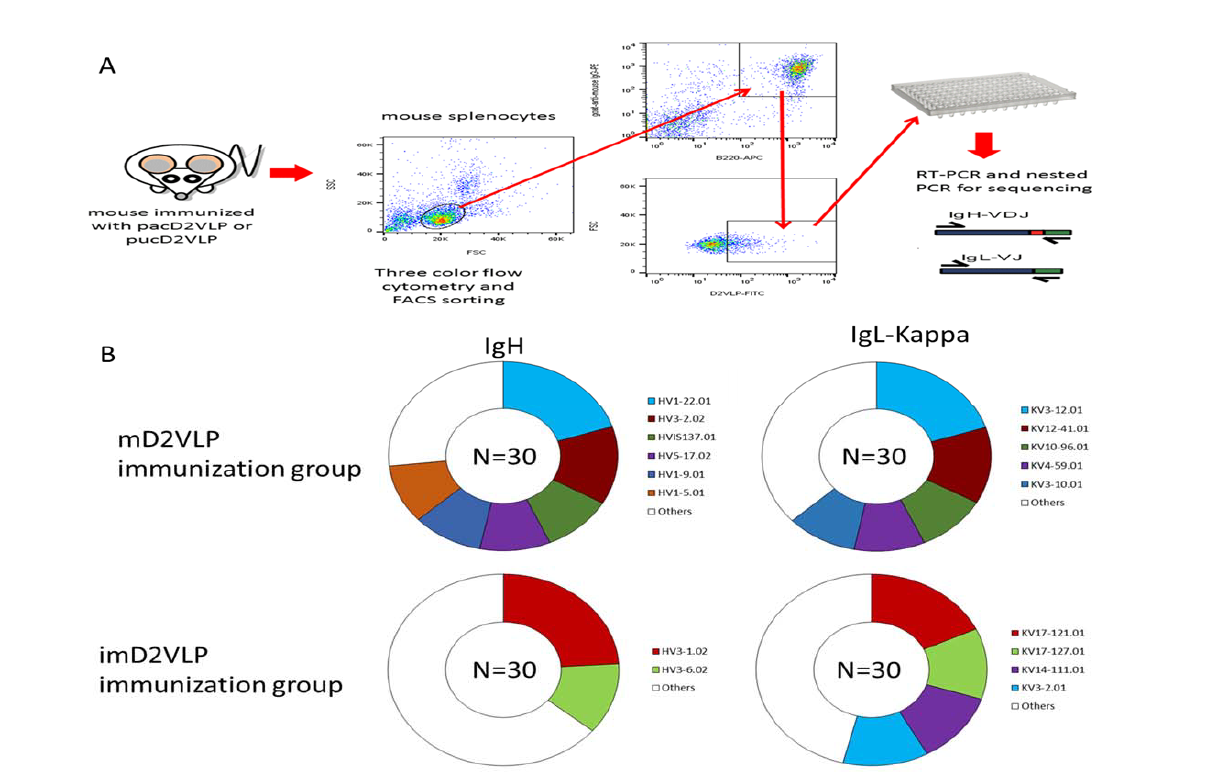
Overview of the experimental design and DENV-specific B-cell repertoires from the splenocytes obtained from mD2VLP- and imD2VLP-immunized mice. (A) Single DENV(+), B220(+) and IgG1(+)-specific B cell from the spleen of vaccinated mice was sorted into 96-well plates by FACS. The IgH and IgL gene transcripts of each single B cell were amplified by RT-PCR and nested PCR using gene-specific primers. (B) B-cell producing heavy chains and light chains encoded by the same IgH and IgL gene loci were grouped by the same color. The proportions of IgH-or IgL (κ-chain only)-genes are indicated by various colors corresponding to frequencies of the B-cell population of each vaccinated group. Frequencies greater than 10% are shown in various colors, otherwise they are grouped together and shown in white.

The major obstacle of dengue vaccine development is the lack of a suitable small animal model to evaluate vaccine efficacy. Therefore, we decided to test whether the monovalent mD2VLP immunogen could provide protection from a lethal dose challenge of heterologous dengue virus serotypes using suckling mice developed in a previous study^21, 38, 39^. The advantage of using sucking mice to test vaccine efficacy is that the protection from dengue virus challenge can only come from maternal antibodies generated by vaccinated female mice and passively transferred to the their babies. The immunization and challenge schedule is illustrated in Fig. 7A. As shown in Fig. 7B, suckling mice born from mothers vaccinated with mD2VLP were all protected from challenge with all four serotypes of dengue virus.

**Fig 7.**
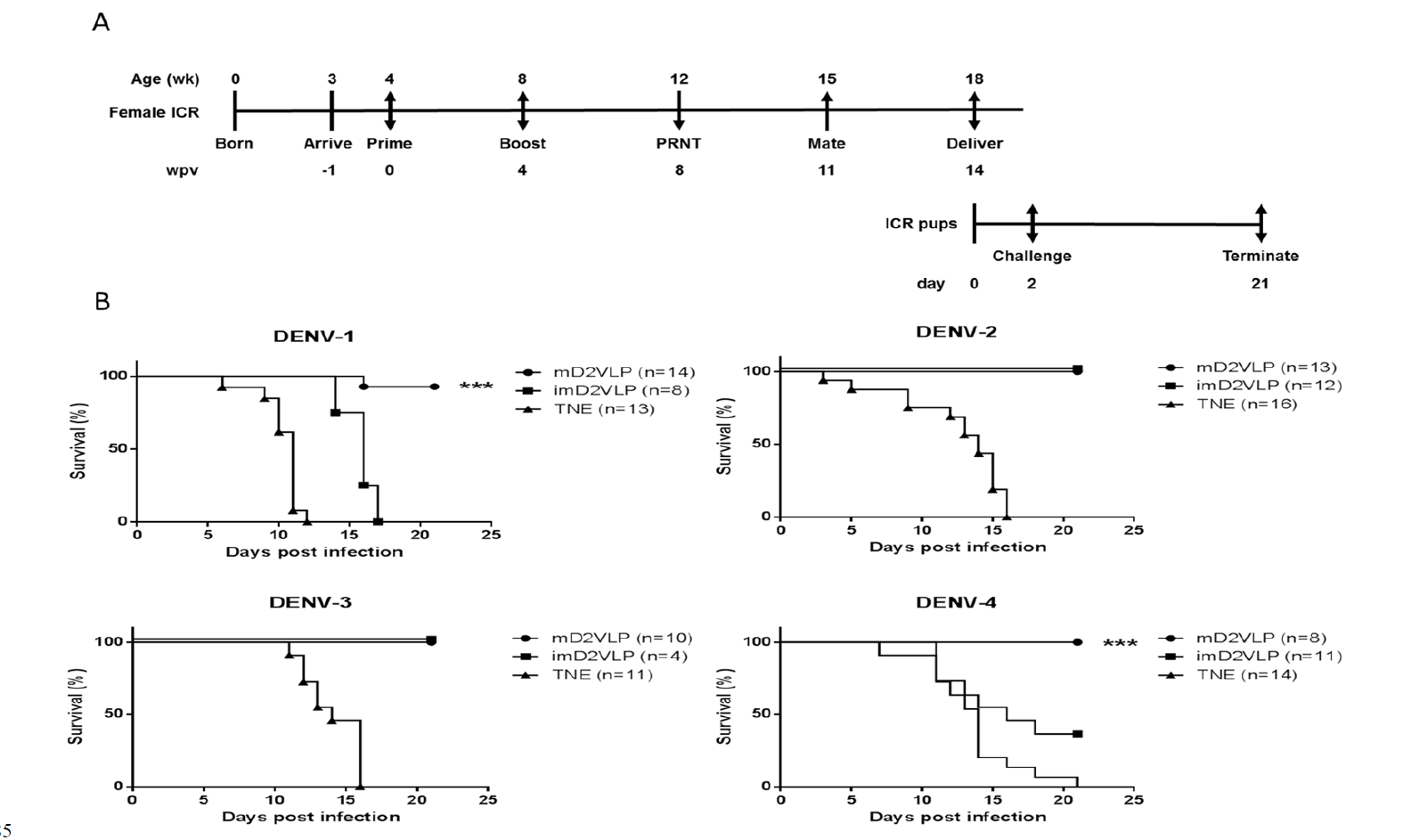
Schematic presentation of the schedule and survival curves for mouse immunization and challenge. (A) Groups of four 4-week-old, female, ICR mice were injected intramuscularly with imD2VLP and mD2VLP at week post vaccination (wpv) 0 and 4 at a dose of 4 μg/100 μL. Mice were bled from the retro-orbital sinus at week 4 following the second injection, and individual mouse serum collected from immunized females 1 week prior to mating was evaluated for the presence of the total IgG titer and the virus neutralization response by ELISA and focus-forming micro-neutralizing assay (FRNT). For the evaluation of passive protection by maternal antibody, ICR pups from the mating of non-immunized males with immunized females 11 weeks post initial vaccination were obtained for viral challenge. Pups from unvaccinated females were used as the challenge control. ICR pups from the designated groups were challenged individually through intracranial route at 2 days after birth with 104 focus-forming unit (FFU) which were equivalent 141, 61, 11, 1000 times of 50% lethal doses (LD50) of DENV-1, to DENV-4, respectively. The percent survival of the mice was evaluated daily for up to 21 days. (B) Survival curve of pups delivered from the female mice receiving mD2VLP, imD2VLP monovalent vaccine or TNE control, then challenge with DENV-1 to 4 after birth. The in vivo protective efficacy of DENV-2 monovalent vaccine is maturity-dependent. N in parentheses indicated the numbers of pups of each group. Kaplan-Meier survival curves were analyzed by the log-rank test. ***, *P* <0.001.

How VLP immunogens mimic their viral counterparts structurally and how the neutralizing epitopes are preserved on the VLP surface are of significant interest in the development of good vaccine candidates. Recent studies suggest that potent human neutralizing antibodies with broad reactivity across dengue serocomplex can be generated from dengue patients, particularly after secondary infection^18, 19, 33^. Our findings here suggested that mD2VLP, with a scaffolding of multiple E-protein dimers similar to that of virion, is capable of inducing such CR antibodies with broad neutralizing activity. In addition, mD2VLP-vaccinated mice produce antisera predominantly composed of about 48% of DM25-3-like antibodies (Fig S11). The packing of E dimers on VLPs provides a unique surface-accessible structure with increased epitope accessibility. Usually these epitopes are cryptic in mature virions maintained at 28oC and in a neutral-pH environment^29^. By superimposing the E-dimer-dependent quaternary epitopes^19, 30, 31^ onto the structure of mD2VLP, the binding footprints of these antibodies were highly overlapping and formed a “neutralization-sensitive hotspot” on mD2VLP (Fig. S12). Thus, the underlying mechanisms of why mD2VLPs are able to stimulate an elevated and broader immune response are based on the following: (1) epitope accessibility exposed at the grooves within E-protein dimers, which govern the generation of neutralizing antibodies; (2) removal of decoy epitopes presented on prM-containing structures found in imD2VLPs; and (3) inter-dimeric epitope accessibility due to the 5-fold and 3-fold openings of the E protein arrangement of T=1. Other mechanisms cannot be excluded such as reducing the production of MAb E53-like antibodies, which have a preference of binding to spikes on noninfectious, immature flavivirions by engaging the highly conserved fusion loop that has limited solvent exposure of the epitope on mature virions^40^. However, studies of vesicular stomatitis virus demonstrated that *in vivo* protection depended on minimum serum antibody concentration, regardless of immunoglobulin subclass, avidity or in vitro neutralization activity^41^. Regardless of the mechanisms involved in NtAb generation and protection from viral challenges, we have found that a novel mature form of monovalent VLP from dengue virus serotype 2 is efficient in inducing elevated and broadly neutralizing antibodies, which makes it an excellent model system for use in exploring the potential of a universal dengue vaccine in humans. Our unique findings in this study showed that a mature, monovalent dengue 2 VLP with grooves within the E protein dimeric molecules on the surface of particles induced highly protective, NtAbs against heterologous dengue viruses from all four serotypes. These characteristics strongly suggested this monovalent VLP as a universal dengue vaccine design without the requirement for tetravalent immunogens. The strategy here may also provide a new direction for the development of other flavivirus vaccines, including Zika virus.

## Acknowledgments

We thank Dr. Felix Rey for commenting on the structure of dengue VLP and Ann Hunt for English editing.

## Author contributions

WFS performed the characterization of D2VLP including Western blotting, ELISA, site-directed mutagenesis and mouse immunization. JHL conducted the Pitou and solvent accessibility analysis. MYL, DYC, HCW and JUG produced and characterized the NS1-splenocytes fused B cell lines, monoclonal antibody and single B-cell repertoire characterization. WFS and YCW conducted VLP purification, mouse immunization experiments and neutralization assays. JUG and CHH prepared the anti-M protein mouse polyclonal serum. JJL and YLL prepared the parental and domain III-replaced recombinant DENV-2. SRC and SRW prepared the cryo-EM samples, processed the cryo-EM images, fitted the protein structures, and solved the cryo-EM structure. MTW, GJC and WFS mapped the 2H2 binding epitopes. SRW, GJC and DYC supervised the studies and contributed to the writing of the paper.

